# Weight Loss in Response to Food Deprivation Predicts the Extent of Diet-Induced Obesity in C57BL/6J Mice

**DOI:** 10.1101/004283

**Authors:** Matthew J. Peloquin, Dave Bridges

## Abstract

Inbred C57BL/6J mice have been used to study diet-induced obesity and the detrimental physiological effects associated with it. Little is understood about predictive factors that predispose an animal to weight gain. To address this, mice were fed a high fat diet, control diet or normal chow diet. Several measurements including pre-diet serum hormone levels and pre-diet body weight were analyzed, but these had limited predictive value regarding weight gain. However, baseline measurements of weight loss in response to food deprivation showed a strong negative correlation with high fat diet-induced weight gain. These data suggest that fasting-induced weight loss in adolescent mice is a useful predictor of diet-induced weight gain.

## Introduction

Obesity is a major global health concern with an estimated 1.4 billion overweight and 500 million obese individuals worldwide [1]. Obesity has a complex etiology including both genetic and environmental inputs. It has been estimated that between 40-70% of obesity is heritable [2]. The non-heritable component of obesity and the factors that modulate it are harder to estimate independently due to variations in diet, exercise and other factors.

Identifying at risk populations of patients and determining how to prioritize limited health care spending is a major public health issue. However, outside of genetic tests for monogenic obesity disorders, there are few good diagnostic criteria for early interventions for at-risk populations. Furthermore, the mechanisms that cause the variable susceptibility to diet-induced obesity are not well understood. Previous work has suggested a variety of factors are predictive of weight gain in human populations, including birth weight [3, 4], adolescent weight [5], leptin [6, 7], and binge eating [5, 8] but these are often difficult to separate from other genetic and socio-economic factors in human populations.

Mouse models of obesity have been important to our understanding of the molecular mechanisms underlying obesity by allowing investigators to control the genetics and environment of animals with very high precision. Inbred C57BL/6J mice are highly genetically similar and are rigorously maintained to reduce genetic drift [9, 10]. Previous studies have identified variable responses to weight gain in inbred C57BL/6J mice and suggested that this was established early in life, shortly after weaning [11–13]. Also, work with rodent models of diet-induced obesity have described a set point in which animals defend their homeostatic body weight [14, 15]. In this study we examine body weight defense, as measured by weight reductions during fasting, to test whether set point maintenance correlates with weight gain.

To test predictive variables that denote susceptibility in an inbred mouse strain we performed diet and genetic induced obesity studies on inbred mice in animal facilities at two sites. Animals were fed either an obesogenic high fat diet (HFD) or one of two control diets (CD or normal chow diet; NCD, see Table 1) and we examined both changes in their physiology and prospective determinants of weight gain.

**Table 1:** Description of Normal Chow Diet, Control Diet and High Fat Diet used during the course of the study. Carbohydrates are sub grouped into sucrose and starch/maltodextrose. Due to seasonal variations in the composition of the Normal Chow Diet, these are best estimates as provided by the vendor.

| Food | Normal Chow | Control Diet | High Fat Diet |
| --- | --- | --- | --- |
| Fat (%) | 17 | 10 | 45 |
| Protein (%) | 29 | 20 | 20 |
| Sucrose (%) | 7 | 17 | 17 |
| Starch/Maltodextran (%) | 47 | 52 | 18 |
| Calories per gram | 3.0 | 3.85 | 4.73 |
| Type | Chow | Synthetic | Synthetic |

## Materials and Methods

### Materials

Male C57BL/6J mice (stock number 000664) and *ob/ob (Lep^ob/ob^*) mice (000632 for C57BL/6J and 0004824 for BTBR background) were ordered from The Jackson Laboratory (Bar Harbor, ME) and received at 8 weeks of age. NCD (8640 Teklad Rodent Diet) was provided by the University of Tennessee Health Science Center Laboratory Animal Care Unit (Memphis, TN) and the University of Michigan Animal Care Facility (Ann Arbor, MI). HFD (D12451) and CD (D12450H) were purchased from Research Diets (New Brunswick, NJ) and stored at 4°C until use. The composition of these diets are presented in Table 1. Note that there are seasonal variations in the composition of the Teklad diet, and therefore these numbers are reasonable estimates on a batch to batch basis provided by the vendor. Blood glucose levels were measured using an OneTouch Ultra 2 Glucometer and OneTouch Ultra Test Strips. All animal procedures were approved by the Animal Care and Use Committee at UTHSC, and the University Committee on Care and Use of Animals UM.

### Animal Housing

Experimental mice from cohorts 1 and 2 were housed at the University of Michigan Animal Care Facility (Ann Arbor, MI). Experimental mice from cohorts 3 and 4 were housed at the University of Tennessee Health Science Center Laboratory Care Unit (Memphis, TN). Mice in diet groups NCD and CD were housed 5 mice per cage, while mice in the HFD group were housed 4 mice to a cage. All mice were kept on a 12/12 light-dark cycle for the duration of the study. Mice being fed NCD and HFD were given 300g of food every 2 weeks, while mice fed CD received 400g. Cage-level food consumption and individual body weights were measured at 2-week intervals at approximately ZT11, at which point the appropriate food was replenished back to the original amount. For the ∼215 day old mouse cohort shown in Figure 2A, these mice were maintained on NCD for a longer duration than other cohorts, before fasting responses were measured.

**Figure 1:**
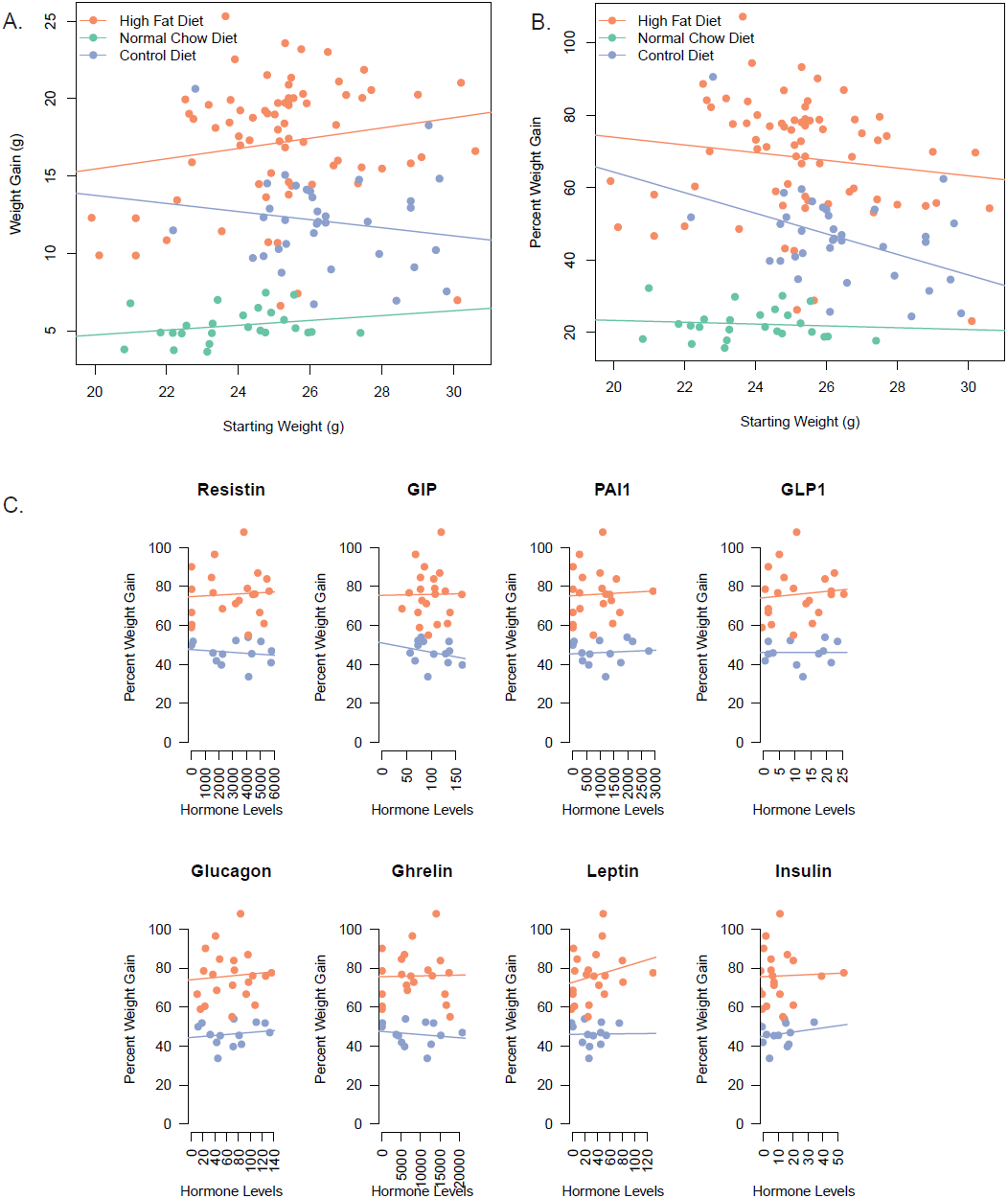
Pre-diet weight and hormone levels have no predictive value for high fat diet-induced weight gain. A) Pre-diet weight of mice compared to weight gain (A) or percent of weight gained (B) while on the diet. C) Fasted hormone levels in serum prior to diet compared to percent weight gain on the diets (also see Table 2).

**Figure 2:**
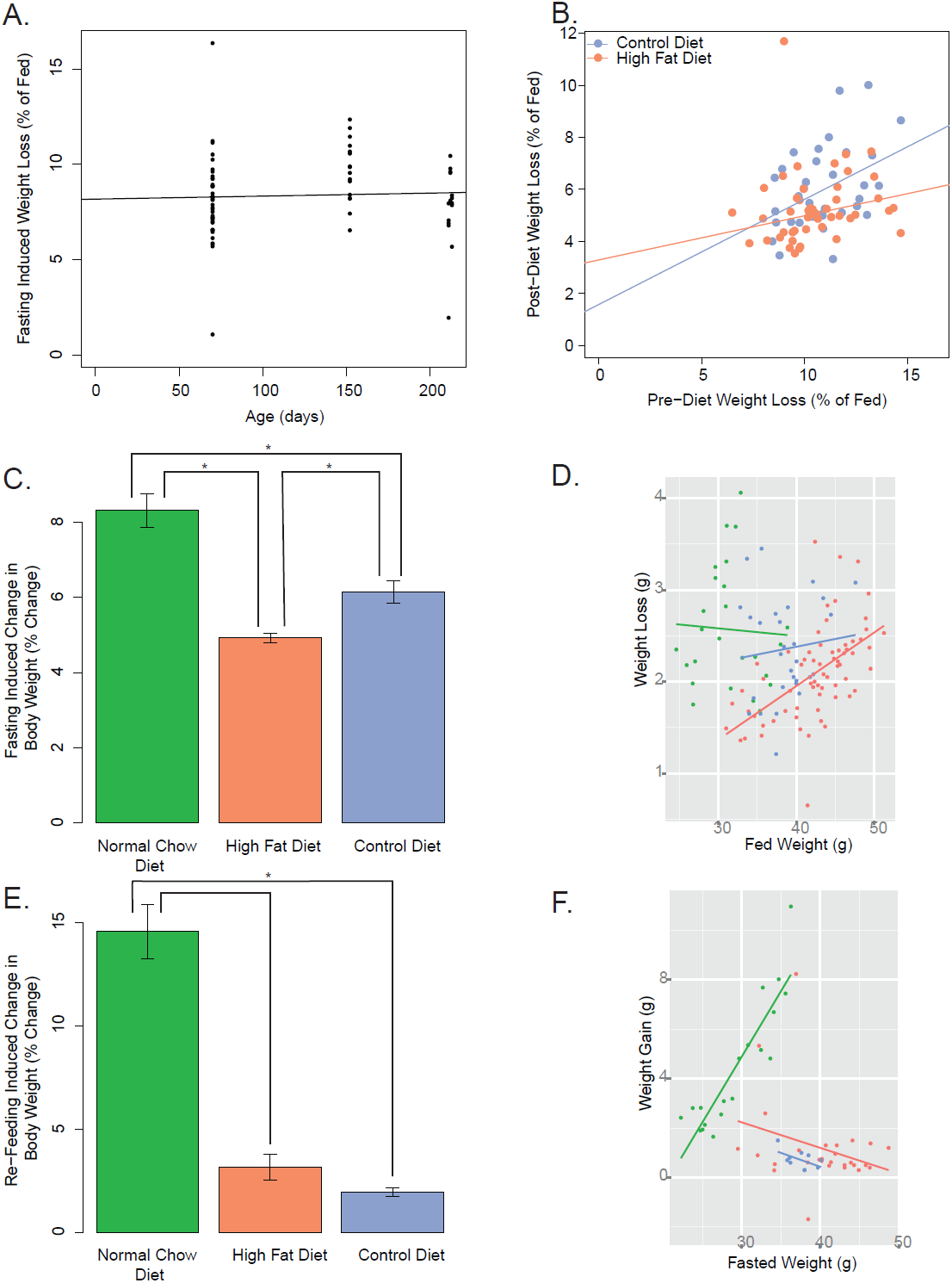
Effects of Dietary Treatments on Fasting Responses. A) Fasting-induced weight loss unaffected by ageing within mice up to ∼200 days. B) Pre-diet weight loss percent shows no significant correlation to post-diet percent of weight loss in HFD and CD fed mice. C) Change in percent of body weight due to fasting for 16 hours for each of the diets. D) Effects of body weight on percent weight loss for each group. E) Change in body weight percent after 6 hours of re-feeding, following the 16 hour fast. F) Effects of body weight on percent weight re-gain for each group. Asterisk indicates a significant difference between groups by Wilcoxon Test after a significant Kruskal-Wallis test (p<0.001).

### Fasting Response and Tissue Collection

Prior to starting the experimental diets, mice were weighed and fasted for 16 hours from ZT11 to ZT4 in clean cages with unrestricted access to water. Following the fast, blood glucose levels, weight measurements were taken and blood was drawn. Following the completion of the 12-week experimental diet treatment, the same fasting protocol was repeated. For re-feeding experiments, the indicated food was re-administered to fasted animals for 6 hours *ad libitum* along with water.

### Hormones and Glucose Measurements

Serum hormone levels were measured using a Bio-Plex Pro Mouse Diabetes Panel 8-Plex kit (BioRad catalog# 171-F7001M) on a Luminex 100/200 plate reader with xPONENT software. Bio-Plex multiplex assay was conducted as described by the manufacturer’s instructions.

Blood glucose levels were taken from all experimental mice after 16 hours of fasting pre-diet and post-diet, while re-fed blood glucose levels were taken for approximately half of the mice. Blood was extracted from the retro orbital vein using a micro hematocrit capillary tube, then placed on ice for 20 minutes to clot, followed by a 20 minute spin at 2000g and storage of the serum at −80C. Glucose was measured from whole blood using an OneTouch Ultra 2 Glucometer.

#### Statistics

Statistical significance for this study was determined with a p/q value of less than 0.05. All statistical analyses were performed using R version 3.1.0[16]. Alterations in food intake were examined using mixed effects linear modeling using lme4 (version 1.0-6 [17]). To determine the effect of diet and time on the diet, we generated a mixed linear model containing the diet type and week as fixed effects and the cage as the random effect. We compared this to models without the week factor and performed a F-test comparing these models. Similarly, to test the effects of diet we compared to a model without the diet type term. To examine the effects of specific differences from HFD fed animals we performed a Dunnett’s test on the mixed effects models using the multcomp package (version 1.3-2 [18]).

Correlations were determined using Spearman’s Rho after testing whether the covariates were normally distributed (Shapiro-Wilk test p<0.05). For potential correlates of weight gain, p-values were adjusted for multiple observations by the method of Benjamini and Hochberg [19]. When a difference or correlation was observed to be significant (or not to be significant) over multiple combined cohorts of mice we also examined each of these cohorts separately. If the combined trend agreed with all of these cohorts independently, we used the combined groups with its increased statistical power. If there was not agreement in all cohorts we do not report this effect as significant, even if the combined analysis suggested it was.

For examining effects of the three dietary treatments, we first performed an ANOVA or Kruskal-Wallis test, depending on whether the data were normally distributed (p>0.05 by Shapiro-Wilk Test). If that was significant, a Tukey post-hoc test or pairwise Wilcoxon Rank Sum Tests were used. All raw data and reproducible analysis code for this manuscript are available at https://github.com/BridgesLab/PredictorsDietInducedObesity.

## Results and Discussion

### Characterization of Effects of Diets on Weight Gain

To test the effects of HFD on weight gain in inbred mice, we placed independent cohorts of 10-week old C57BL/6J male mice on either a NCD, CD or HFD for 12 weeks (see Supplementary Figure 1A). Often, mice are maintained on NCD, but due to the substantial differences in chemical make up of this diet and synthetic diets we also tested a control diet. The CD had less fat (10% vs. 40% of calories from fat compared to the HFD) and also had identical protein and sucrose content (Table 1). We examined the effects of weight gain over four separate cohorts of mice, and found that the HFD-fed mice gained substantially more weight than the CD or NCD mice, but also that the CD mice gained substantially more weight than the NCD mice (Supplementary Figure 1B). HFD fed mice weighed significantly more at all time points during the 12-week diet treatment compared to the NCD fed mice in cohorts 1 and 2, as well as CD fed mice in cohorts 3 and 4 (Supplementary Figure 1B–C).

To probe the effects of the diet on food intake, we measured the amount of food consumed by each cage of mice on a bi-weekly basis. We found that food intake on both a per gram basis (Supplementary Figure 1D, p=0.0033) and a caloric basis (Supplementary Figure 1E, p=0.0038) decreased over time even as the mice gained weight. There was a significant effect of diet (p<0.005 for both caloric and absolute food intake by F-Test) on the amount of food intake. Specifically, we observed that the CD fed animals consumed 2.0 kcal/mouse/day less food than HFD fed animals (corresponding to 1.1 g/mouse/day more; p<0.005 for each comparison). The NCD fed animals also ate more grams of food (0.48 g/mouse/day p=2.8 × 10^−5^), but only slightly less calories than the HFD fed animals (p=0.18).

To test whether synthetic CD generated similar metabolic changes as NCD to HFD comparisons, we examined serum hormone levels of key obesity related factors in both the fasted and re-fed conditions (Supplementary Figure 2A) and blood glucose levels (Supplementary Figure 2B). Significant differences of were detected between several hormones (resistin, total ghrelin, GLP-1 and leptin) as well as fasting glucose levels between these diets. These are consistent with previous reports of HFD induced changes relative to NCD diets for resistin and leptin [20, 21]. Futhermore, the fasting/re-feeding responses (decreases in total ghrelin, glucagon, GLP-1 and increases with insulin) are consistent with previous data, supportive of a normal fasting response [22–25]. Based on these data, the hormonal responses in these 16h fasted mice appear normal.

### Predictors of Weight Gain

To understand the variance in body weight that occurs within an inbred mouse line, we tested several potential factors for their ability to predict weight gain. We first investigated if pre-diet body weight had any correlation with eventual weight gain (Figure 1A and B). We observed no significant correlation between initial pre-diet body weights and weight gain post-diet in both in terms of absolute weight gain (p=0.089, Rho^2^=0.12) and percent weight gain (p=0.89, Rho^2^=0.0008) for HFD fed animals. We also did not observe any correlation for between initial weight and weight gain for NCD or CD fed animals.

We next examined serum collected from mice before they were placed on the diets, to test the hypothesis that pre-diet serum hormone levels are associated with weight gain in HFD fed mice. We observed no significant correlation between pre-diet hormone levels and either percent or absolute weight gain in both HFD and CD fed mice (Figure 1C and Table 2). Collectively, these data suggest that pre-diet body weight and common metabolic hormone levels are not predictive of weight gain in male C57BL/6J mice.

**Table 2:**
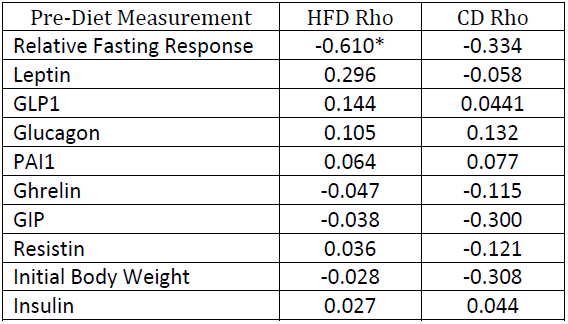
Correlation between Pre-Diet Measurements and Percent Weight Gain on a Control or High Fat Diet. Spearman’s Rho coefficient is shown for each predictor. Asterisk indicates q<0.05 after correcting for multiple observations.

| Pre-Diet Measurement | HFD Rho | CD Rho |
| --- | --- | --- |
| Relative Fasting Response | -0.610* | -0.334 |
| Leptin | 0.296 | -0.058 |
| GLP1 | 0.144 | 0.0441 |
| Glucagon | 0.105 | 0.132 |
| PAI1 | 0.064 | 0.077 |
| Ghrelin | -0.047 | -0.115 |
| GIP | -0.038 | -0.300 |
| Resistin | 0.036 | -0.121 |
| Initial Body Weight | -0.028 | -0.308 |
| Insulin | 0.027 | 0.044 |

One hypothesis is that one dominant mouse may affect the weights of other mice in its cage. To test this we looked at the 5 heaviest from our data, existing in 5 distinct cages. Those cages contained 20 mice in total. The other 15 mice in these cages (excluding the heavy ones) weighed on average slightly more than the average of all other mice analyzed. Since these mice did not weigh significantly lower (p=0.392), they do not support the hypothesis that one heavy, dominant mouse drives the weights of its cage-mates to be lower (Supplementary Figure 4). By the same token, the presence of a larger mouse does not make the other mice in that cage mice significantly heavier.

### Fasting Response Predicts Weight Gain

Another factor that we examined was the effect of food deprivation on diet induced obesity. To do this, we deprived mice of food for 16h both before the dietary intervention, and after it. Fasting responses, as measured by either absolute or percent reductions in body weight were stable within NCD fed male C57BL/6J mice over time (Figure 2A, Rho^2^<0.02, p>0.26), and only weakly correlated with the pre-diet weight fasting responses within mice in either relative or absolute terms (Figure 2B, Rho^2^<0.12, p<0.019 for HFD and Rho^2^<0.153, p<0.031 for CD).

As shown in Figure 2C, fasting-induced weight loss was significantly higher in NCD mice than in the HFD or CD mice (p<0.0005) after the dietary treatment. CD-fed mice also had a stronger fasting response than HFD mice (p=0.00018). We tested whether the post-diet body weight could explain these differences in fasting and re-feeding responses. Globally, there was a no significant correlation between body weight and the absolute weight lost during the 16 fast (p=0.881). When we separated the mice by dietary treatment we observed a significant positive correlation between body weight and absolute weight loss in the HFD treated mice only (Rho^2^ = 0.404, p=7.5 × 10^−11^, Figure 2D). If we examine percent weight loss rather than absolute weight loss, there is no correlation between weight loss and body weight in the HFD fed animals (p=0.22).

When the fasted mice were re-fed for 6 hours, NCD fed mice gained significantly more body weight than either HFD or CD fed mice (p<3 × 10^−5^, Figure 2E). There was no statistically significant difference in re-feeding induced weight gain between CD and HFD mice (p=0.7). In the case of NCD (Rho^2^ >0.65, p<4.2 × 10^−5^) but not the two synthetic diets (HFD or CD), there was a strong positive correlation between body weight and their re-feeding induced absolute or percent weight gain over those 6 hours (Figures 2F). These data suggest that response to re-feeding is strongly altered by dietary type (synthetic versus chow) but is largely independent of their body weights.

For leptin mutant *ob/ob* mice, we observed inconsistent results between backgrounds. We examined the fasting response between genetically obese mice on either a BTBR or C57BL/6J genetic background. *ob/ob* mice are leptin deficient due to a single nucleotide polymorphism in the leptin gene. This results in mice which eat the same type of food (NCD), but substantially more of it, leading to obesity [26–29]. The effects of this mutation on body weight and fasting glucose in these strains are shown in Supplementary Figures 3A-B). For C57BL/6J-*ob/ob* mice, we observed a significant increase in fasting induced weight loss relative to control mice, opposite to what we observed for HFD-fed mice (Supplementary Figure 5). However, for BTBR-*ob/ob* mice, we observed an increase in the percent weight loss. These data suggest that background differences may play a role in fasting response in the absence of leptin.

We then tested whether the pre-diet fasting response is predictive of eventual weight gain during the course of the diet for the two synthetic diets. Due to the dramatically different fasting responses observed in NCD mice compared with CD mice (See Figure 2C and E) we only performed these studies comparing the more chemically comparable CD and HFD diets. Both HFD and CD fed mice exhibited less weight gain when the pre-diet fasting response was elevated (Figure 3). We found a strong negative correlation between percent weight gain and relative fasting response (Rho=-0.61, Rho^2^ =0.37, p=6.6 × 10^−6^). For CD fed mice the same trend was present, though the correlation did not quite reach statistical significance (Rho=-0.334, Rho^2^=0.11, p=0.054). These trends also reached statistical significance for all the combinations tested between both relative and absolute fasting responses and relative and absolute weight gain during HFD (p<0.03 and Rho<-0.33 for all comparisons). These data show that mice which resisted weight loss during the pre-diet 16 hour fast were far more susceptible to weight gain while on the experimental diet, and that this factor predicted weight gain much more comprehensively than hormonal factors, or baseline weight (Table 2 and Figure 3).

**Figure 3:**
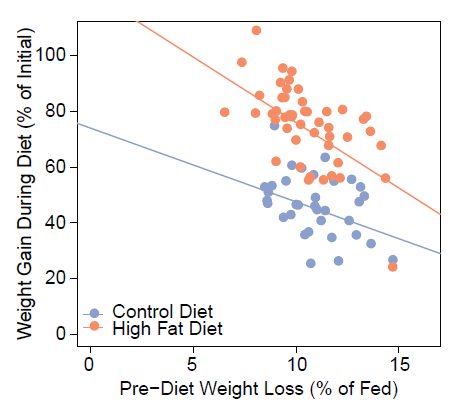
Pre-diet fasting response negatively correlates with weight gain. Correlation between relative fasting induced weight loss at 10 weeks and percent weight gain during the 12 week diet.

## Conclusions

In this study we have described the physiological effects of dietary manipulation in a common inbred strain of laboratory mice. The aim of this study was to control the genetic background, environment and diet of these laboratory animals as closely as possible in order to assess the amount of variability that is not due to genetic differences.

We have observed substantial within-strain variability in the response to HFD and have explored the physiological basis for these differences by examining a variety of pre-diet biomarkers. We did not observe any data supporting the hypothesis that baseline body weight or fasting hormone levels are significantly predictive of weight gain. We did observe a strong predictive effect of body weight responses to food deprivation on diet-induced obesity. Based on these data, the predictive utility of fasting responses is nearly 5-fold greater than that of leptin levels, which had been previously reported to be of some use in predicting weight gain [30]. The negative relationship between fasting responses and weight gain is similar in magnitude to the positive relationship between absolute fat mass and HFD-induced weight gain in mice [12, 13]. It is possible that young mice that have larger fat stores are able to lose more weight in response to fasting than lean mice and that fasting responses and fat mass levels are related. Understanding the relationship between fat mass, fasting responses and predisposition to weight gain will be the focus of future studies.

This study does not attempt to address the underlying molecular mechanisms for these differences in fasting responses but suggests that there is a physiological state established by the time the dietary treatment begins that causes differential weight gain. One possibility is that alterations in their basal metabolic rate cause these differences, as has been proposed in pediatric human populations [31, 32]. The underlying molecular mechanism may yet be some level of *de novo* genetic variation in these mice, or epigenetic modifications that alter sensitivity to dietary factors. This study provides a phenotypic framework to test these molecular hypotheses.

We tested the effects of prolonged food deprivation, as this intervention will activate not only glucagon signaling, but also other catabolic signaling cascades including growth hormone and glucocorticoid [33, 34]. In this way, our protocol for food deprivation is likely to engage all of these pathways to defend body weight in the absence of food. Of note it is interesting that fasting responses themselves are not stable throughout life on a per mouse basis (see Figure 2B). This suggests that either the fasting response is not causative of weight gain directly, or only correlates with predisposition to weight gain during a younger age. These data are consistent with reports that among adult human populations, basal metabolic rate is not reduced in obese individuals [35, 36] while reductions in metabolic rates in pediatric populations are predictive of obesity [31, 32]. Furthermore, we noted that the negative relationship between fasting responses and weight gain is present for both CD and HFD fed animals. This suggests that the fasting response may be part of a general body weight defense mechanism, rather than a phenomena specific to diet-induced obesity, and that this effect may be exacerbated by HFD.

These data support a model where susceptibility to weight gain is at least in part caused by a non-genetic factor which is established early in life. This is consistent with previous studies on these mice which also proposed that susceptibility to weight gain is determined early in life [11–13]. Understanding the mechanistic basis for the relationship between fasting induced weight loss and eventual weight gain may be relevant to providing better individualized care of pediatric populations, since it may help predict susceptibility to weight gain in young children.

## Acknowledgements

The authors would like to thank the Cormier Lab (Department of Pediatrics, UTHSC) for assistance with and use of the Luminex system. We would also like to thank Drs. David Buchner (Case Western Reserve University), Irit Hochberg (RAMBAM Health Care Campus), Shannon M. Reilly (University of Michigan), and Joan Han (University of Tennessee Health Sciences Center) for helpful suggestions and members of the Bridges and Reiter laboratories for insightful discussions.

## Supplementary Figure Legends

**Supplementary Figure 1:**
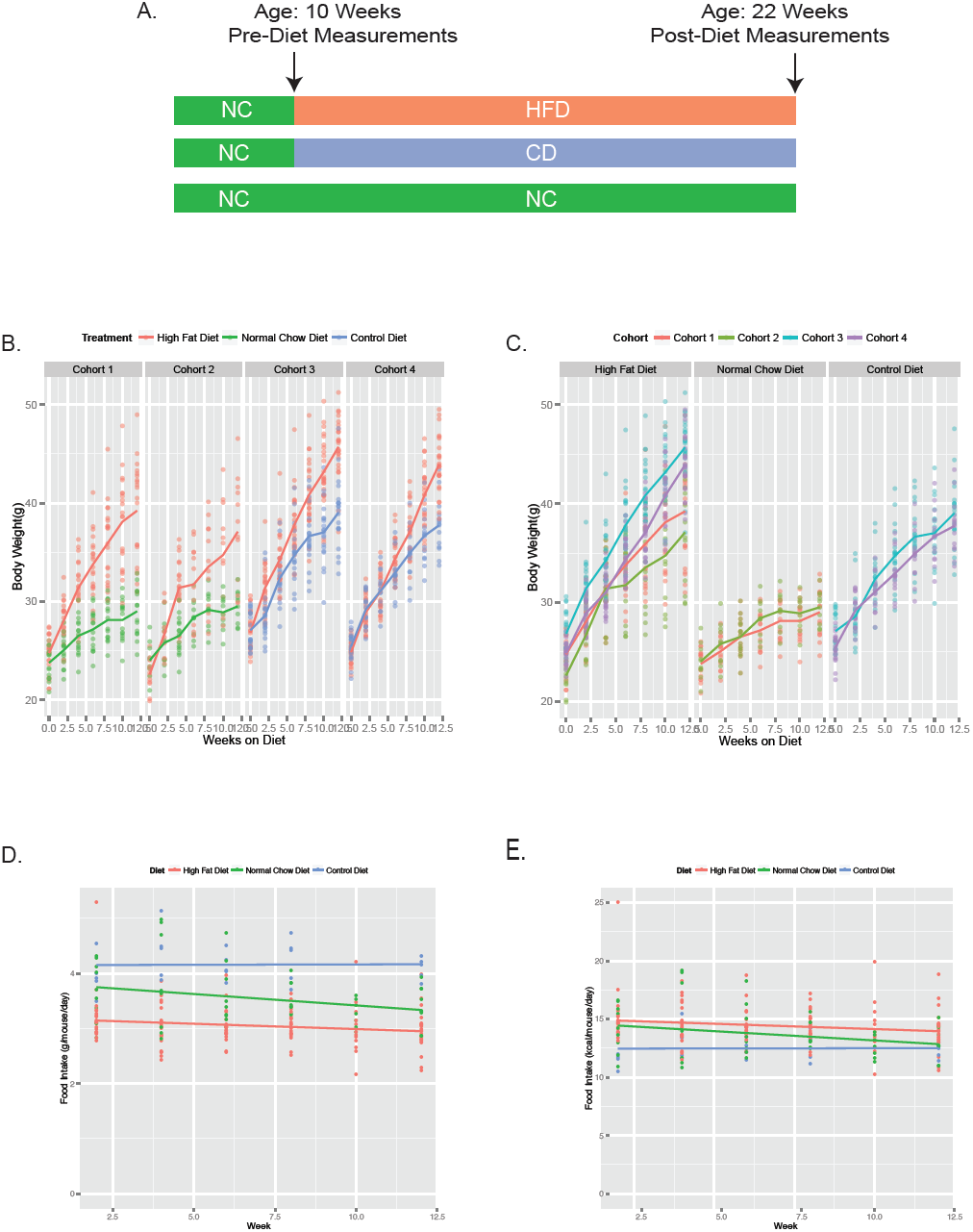
High Fat Diet fed mice gain significantly more weight than Control Diet and Normal Chow diet fed mice. A) Schematic of dietary treatments. Mice were fed NCD from birth to 10 weeks, whereupon the diet was then changed to HFD, CD or remained on NCD for the following 12 weeks. B) Body eights across all studied treatment groups separated by cohorts starting at 10 weeks of age. C) Weight gain across all cohorts treatment groups separated by diet. Food intake per diet over the length of the 12-week diet treatment measured in kcal (D) or grams (E). Food was weighed at the start and conclusion of each 2-week measurement period and each dot represents a single cage at that time point.

**Supplementary Figure 2:**
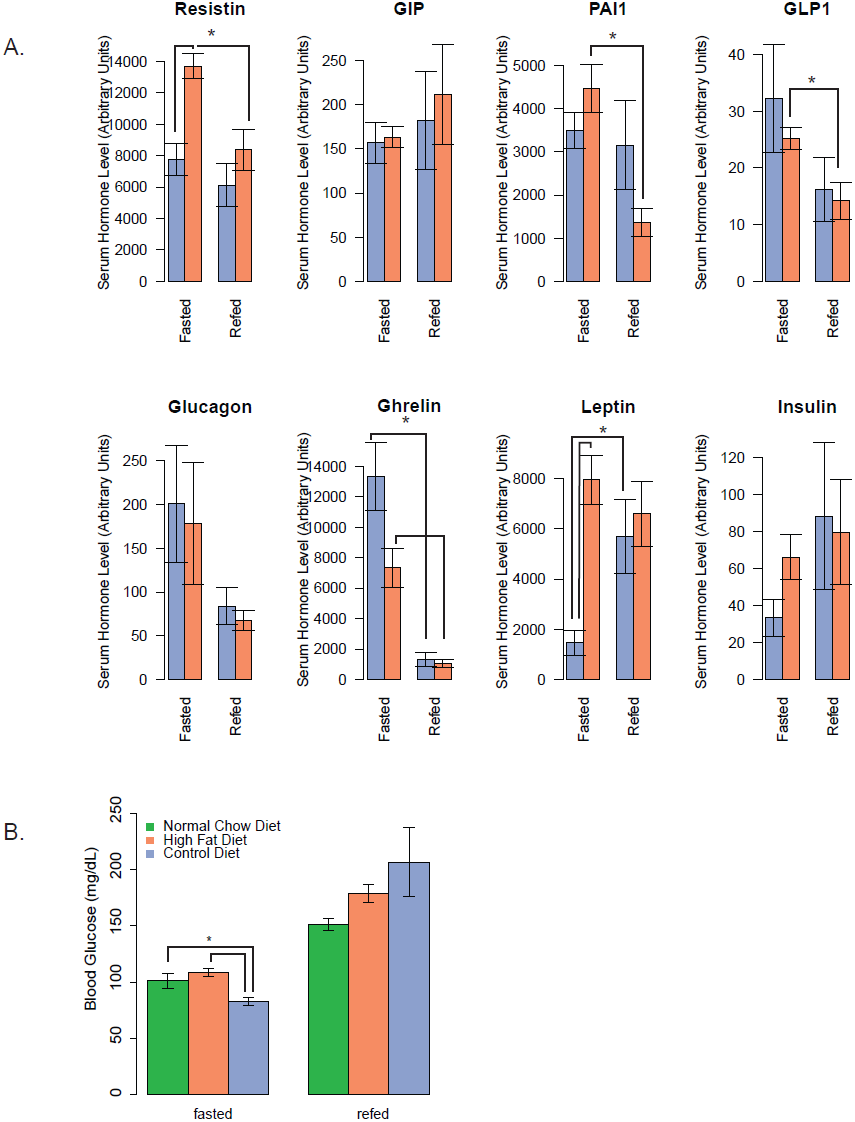
Post-diet hormones hormone levels are similar to previously investigated levels. A) Hormone levels in post-diet serum for HFD and CD mice. Fasted mice were fasted for 16 hours prior to blood collection. Re-Fed mice were given the indicated diet for 6 hours following the 16-hour fast prior to blood collection. Asterisk indicates p<0.05 from pairwise test after a significant ANOVA. B) Blood glucose levels of post-diet fasted and re-fed mice across all treatment groups. Asterisk indicates Tukey test following a significant ANOVA result (B).

**Supplementary Figure 3:**
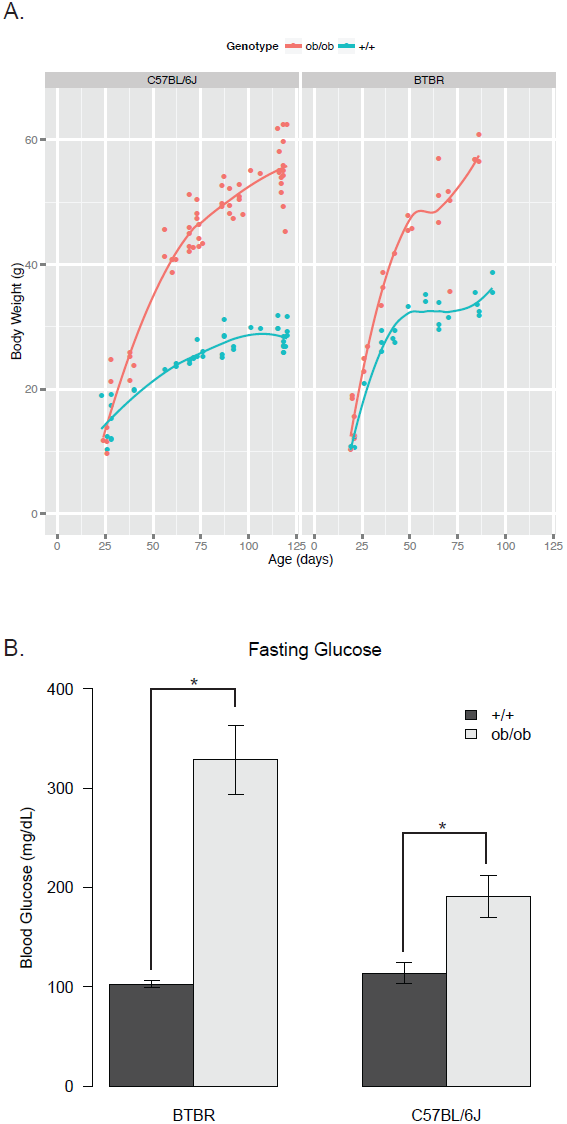
*ob/ob* mice exhibit obesity and hyperglycemia. A) Weights of male *ob/ob* and wild-type littermates on BTBR and C57BL/6J backgrounds. B) Fasting glucose levels between *ob/ob* knockout and wild-type mice. Asterisk indicates p <0.01 via a Wilcoxon rank sum test.

**Supplementary Figure 4:**
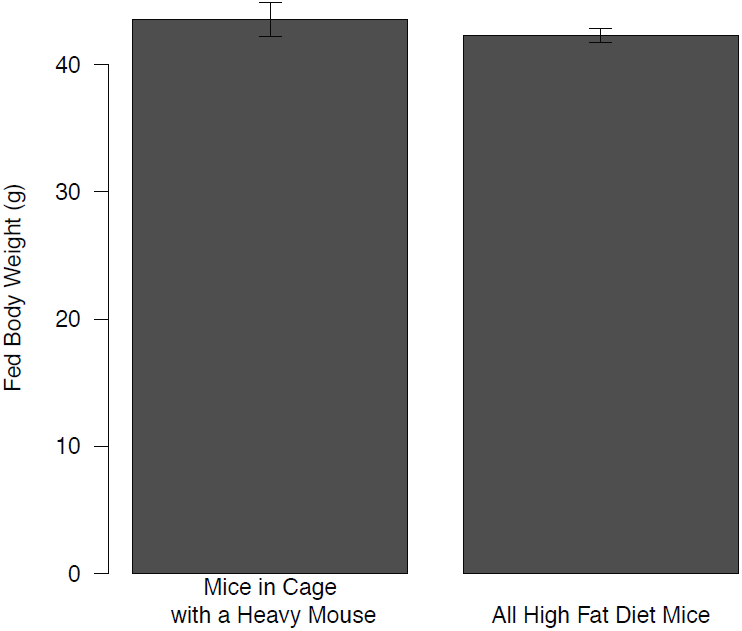
Largest mice did not have a suppressive on the weight of other mice within the same cage. The body weight from the cages containing the 5 heaviest mice in our cohort (with the heavy mouse excluded) was compared to the average of all other mice. No difference was detected between these groups.

**Supplementary Figure 5:**
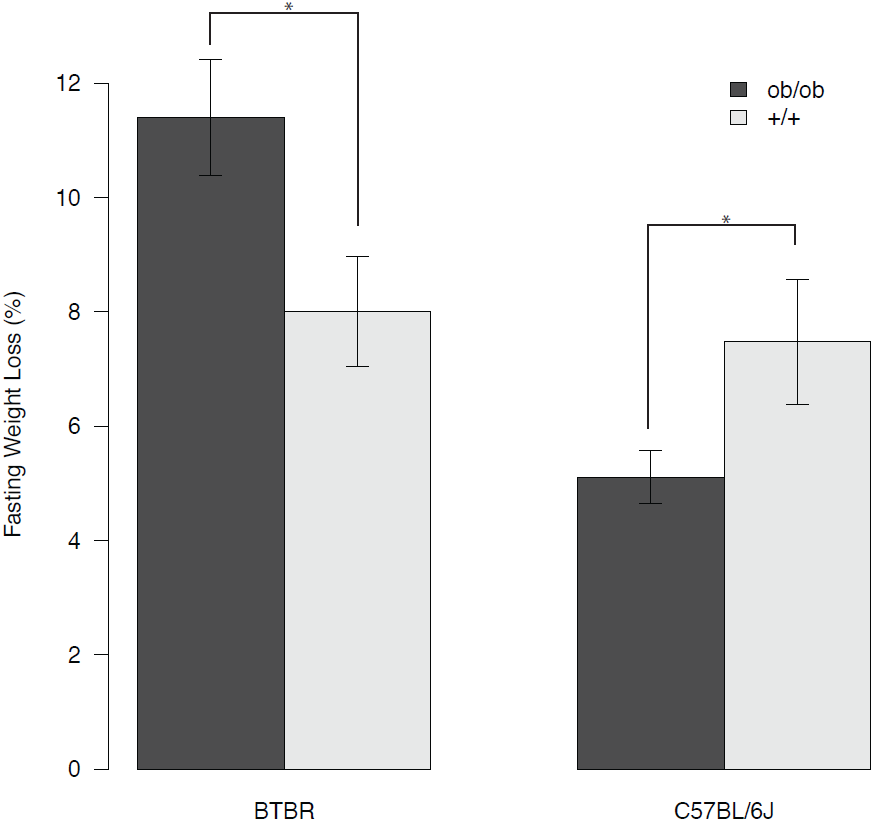
Fasting induced weight loss in *ob/ob* mice. 120 day old *ob/ob* mice were fasted for 16h and the percent weight loss upon fasting was determined. Asterisk indicates p <0.05 via a Wilcoxon rank sum test.

